# Sign inversion in selection on ploidy

**DOI:** 10.1101/2021.10.26.465943

**Authors:** Yevgeniy Raynes, Daniel M. Weinreich

**Affiliations:** Department of Ecology, Evolution, Brown University Box G-W, Providence, RI 02912 USA; Organismal Biology and Center for Computational Molecular Biology, Brown University Box G-W, Providence, RI 02912 USA

## Abstract

Ploidy – the number of homologous chromosome sets in a cell – is remarkably variable across the natural world, yet the evolutionary processes that have resulted in such diversity remain poorly understood. Here we use stochastic agent-based simulations to model ploidy evolution under the influence of indirect selection, i.e., selection mediated solely by statistical associations with fitness-affecting mutations. We find that in non-equilibrium asexual populations, the sign of selection on ploidy can change with population size – a phenomenon we have previously termed sign inversion. In large populations, ploidy dynamics are dominated by indirect effects of selection on beneficial mutations, which favors haploids over diploids. However, as population size declines, selection for beneficial mutations is neutralized by random genetic drift before drift can overwhelm selection against the cost of the deleterious mutational load. As a result, in small populations indirect selection is dominated by the cost of the deleterious load, which favors diploids over haploids. Our work adds to the growing body of evidence challenging established evolutionary theory that population size can affect only the efficiency, but not the sign, of natural selection.

## Introduction

Why do some species have a single set of homologous chromosomes while others have two or more? The evolutionary forces that govern transitions between haploidy, diploidy, and polyploidy and have produced the variability we see today have long been a subject of significant theoretical and empirical interest (reviewed in: Otto & Gerstein, 2008, Gerstein & Sharp, 2021). Theories for the role that natural selection plays in ploidy evolution consider either the immediate effects of ploidy on the phenotype or longer-term consequences for the supply rate of fitness-affecting mutations and the efficacy of selection (Otto & Gerstein, 2008). When the phenotypic effects of ploidy alter the fitness of an individual, ploidy itself may be directly influenced by selection. For example, haploid yeast cells are generally smaller and have a higher surface area to volume ratio than diploid yeast cells, which may afford them higher fitness in nutrient limited conditions (Gerstein & Sharp, 2021, Gerstein et al., 2008, but see Mable, 2001). Ploidy variants may also experience indirect selection acting through statistical associations (i.e., linkage disequilibrium) that develop with fitness-affecting mutations. Indirect selection is particularly effective in asexual populations as recombination can quickly erode linkage disequilibrium.

Most theoretical studies of indirect selection have made progress by considering the effects of selection on deleterious mutations separately from the effects of selection on beneficial mutations. Selection against deleterious mutations can favor diploids over haploids in non-equilibrium populations as diploids can mask the effects of recessive or partially dominant deleterious mutations: i.e., mutations that are fully expressed in haploids but only partially, or not at all, in diploids, where they first appear in heterozygous state (Charlesworth, 1991). In other words, while in haploids such mutations lower fitness by a selection coefficient *s_d_*, in heterozygous diploids their effect on fitness is reduced to *h_d_s_d_*; *h_d_*, known as the dominance coefficient, measures the dominance of the deleterious mutation and is below 1 for any but the fully dominant mutations. However, this advantage in the fitness cost of deleterious mutations may only be a transient one. Once at mutation-selection equilibrium, diploids – with twice as many loci as haploids – end up bearing twice the cost of the deleterious mutational load (Crow & Kimura, 1965, Haldane, 1937). That is, while the equilibrium mean fitness of a haploid populations is 1-*U_d_* (where *U_d_* is the deleterious mutation rate) diploids have an equilibrium mean fitness of 1-2*U_d_* (Crow & Kimura, 1965). Nevertheless, two earlier studies have shown that masking of deleterious mutations could favor diploids at equilibrium in certain limited circumstances. Kondrashov and Crow (Kondrashov & Crow, 1991) found that diploid equilibrium mean fitness could exceed that of haploids under truncation selection and a relatively low *h_d_* (below 0.25). Meanwhile, Perrot et al (Perrot et al., 1991) demonstrated that diploids could invade haploids in populations with random mating between haploids and diploids, high rate of free recombination, and *h_d_* below 0.5; later work confirmed that diploids fail to supplant haploids in populations with rare recombination (Otto & Goldstein, 1992, Otto & Marks, 1996).

Diploids may also be favored because of their increased supply of beneficial mutations, given their 2-fold advantage in beneficial mutation rate. Specifically, if recombination is sufficiently rare diploids may preferentially increase in frequency by hitchhiking (Maynard Smith & Haigh, 1974) along with those beneficial mutations that survive drift and sweep to fixation. However, beneficial mutations with dominance coefficient, *h_b_*, below 1 would also be at least partially masked in diploids, reducing their fitness advantage to *h_b_s_b_* and lowering their fixation probability compared to those in haploids. This advantage in the efficacy of selection could, in certain circumstances, favor haploids. Orr and Otto (Orr & Otto, 1994) compared the rates of adaptive evolution in diploids and haploids. They found that diploids adapt faster than haploids when the rate of adaptation is limited by the supply of beneficial mutations – e.g., when beneficial mutations are rare or population size is small – provided new mutations are at least partially dominant (*h_b_* > 0.5). On the other hand, as long as *h_b_* < 1, diploids evolve slower than haploids when the rate of adaptation is limited not by the supply of beneficial mutations – e.g., in large populations in which such mutations are common – but by the strength of selection for beneficial mutations.

In contrast to most earlier treatments, here we model the evolution of ploidy in non-equilibrium populations with both beneficial and deleterious mutations. Using individual-based simulations we show that, in the absence of recombination, whether ploidy variants are favored or disfavored by indirect selection depends on population size. In small asexual populations, selection against the deleterious mutational load favors diploids and disfavors haploids. In large asexual populations, the cost of the deleterious load is overwhelmed by selection for beneficial mutations, which favors haploids over diploids.

## Methods

We model asexual, clonally reproducing populations of constant size, *N* evolving in discrete, non-overlapping generations. Populations are composed of genetic lineages defined by ploidy state and the number of beneficial and deleterious mutations. Each new beneficial mutation increases an individual’s fitness by a constant *s_b_*; a new deleterious mutation reduces fitness by a constant *s_d_*. We assume that fitness effects are additive and calculate the fitness of a haploid lineage with *x* beneficial and *y* deleterious mutations as w_xy_ = 1+*xs_b_*-*ys_d_*. As in (Orr & Otto, 1994) we also assume that in the absence of recombination fitness-affecting mutations in diploids remain permanently heterozygous (at least on the time scale of our simulations) and calculate the fitness of a diploid lineage with *x* beneficial and *y* deleterious mutations as w_xy_=1+*xh_b_s_b_*-*yh_d_s_d_*, where *h_b_* and *h_d_* are the corresponding dominance coefficients.

At the outset of simulation, a single individual of a particular ploidy (either haploid or diploid) is introduced to a population of *N*-1 individuals of the other ploidy. Both the invading lineage of size 1 and the resident lineage of size *N*-1 bear no fitness-affecting mutations and, thus, have an initial fitness *w* = 1. Every generation the population reproduces according to the Wright-Fisher model (Ewens, 2004) with the size of the lineage carrying *x* beneficial mutations and *y* deleterious mutation in generation *t*+1 randomly sampled from a multinomial distribution with expectation 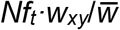 where *N* is the size of the population, *f_t_* is the frequency of the lineage in generation *t, w_xy_* is the fitness of the lineage as defined above and 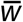 is the average fitness of the population. Upon reproduction, each surviving lineage acquires a random number of beneficial and deleterious mutations drawn from a Poisson distribution with expectation given by, respectively, *cN_i_U_b_* and *cN_i_U_d_*. Here, *c* is the ploidy of the lineage (i.e., number of chromosome copies), *N_i_* is the size of the lineage, *U_b_* is the per-individual beneficial mutation rate, and *U_d_* is the per-individual deleterious mutation rate. Diploids, thus, have a 2-fold higher mutation rate than haploids. For simulations in Fig. 2C we lowered the diploid mutation rate to that of haploids by setting *c* = 1 for both diploid and haploid lineages.

Simulations end when the invader either fixes (reaches a frequency of 1.0) or is eliminated from the population. We assess the fixation probably (*P_fix_*) as the frequency with which the invader supplants the resident across the replicate runs of the simulation. Throughout, we focus on *NP_fix_* - the fixation probability normalized by the neutral expectation, 1/*N*. To ascertain whether an invading ploidy variant is favored or disfavored by selection we compare its *NP_fix_* to that expected of a neutral mutation, i.e., *NP_fix_* = 1 (e.g., Raynes et al., 2018).

Simulation code was written in Julia 1.2 and is available at https://github.com/yraynes/Ploidy.

## Results

We first examined the fixation probability of haploid invaders in diploid populations. Since deleterious mutations in nature are predominantly recessive (e.g., Simmons & Crow, 1977, Wilkie, 1994, Peters et al., 2003), we set the dominance coefficient of deleterious mutations in our simulations to be *h_d_* = 0.0. Meanwhile, we varied the dominance coefficient of new beneficial mutations from fully recessive (*h_b_* = 0.0) to fully dominant (*h_b_* = 1.0). Strikingly, we found that the sign of selection on haploids can change with population size (Fig 1A). For all values of *h_b_* < 1.0 studied, haploids are favored by selection (have *NP_fix_* > 1) in large populations but become disfavored (have *NP_fix_* < 1) as population size decreases below a critical size we call *N_crit_*. We have previously designated this *N*-dependent switch in the sign of selection *sign inversion* (Raynes et al., 2018, Raynes et al., 2021). Fig. S1 confirms that haploids are susceptible to sign inversion across a wide – at least over several orders of magnitude – range of mutation rates.

**Figure 1:**
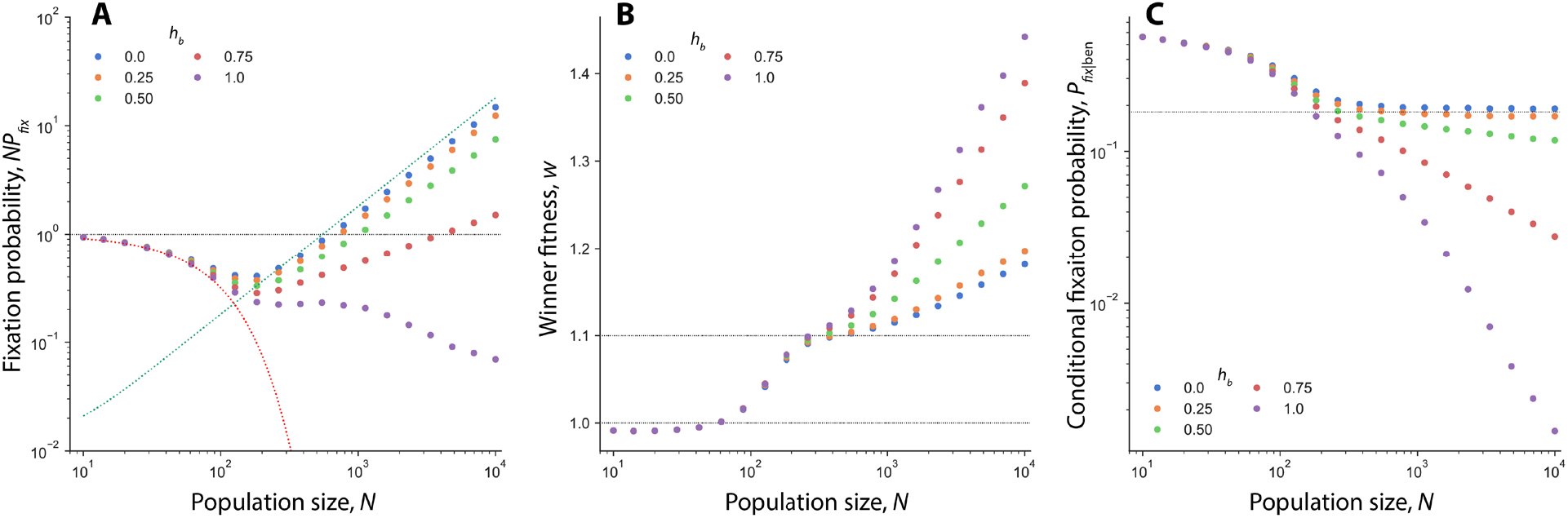
Sign inversion in selection on haploids. **A)** Normalized fixation probability (*NP_fix_*) of haploid invaders (circles) in diploid populations at different *h_b_*. Red dashed line: *P_fix_*(*s* = -*U_d_, N*); Teal dashed line: 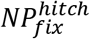. Horizontal black line shows the normalized fixation probability of a neutral mutation (*NP_fix_* = 1). **B)** Average fitness of winning haploid populations, *w*. Horizontal black lines indicate *w* = 1 and *w* = 1 + *s_b_* = 1.1 **C)** Conditional fixation probability calculated across simulations in which haploid invaders managed to acquire at least one beneficial mutation. Horizontal black line shows expected *P_fix_*(*s_b_, N*). In all panels, parameter values are as follows: *s_b_* = 0.1, *s_d_* = −0.1, *U_d_* = 0.01, *U_b_* = 0.001. All simulation results averaged across at least 10^7^ replicates.

Based on our earlier work on sign inversion (Raynes et al., 2021, Raynes et al., 2018) we assumed that in small populations haploid *NP_fix_* would be reduced below the neutral expectation by selection against the deleterious load. To test this expectation, we compared haploid *NP_fix_* in simulations to that predicted by the fitness cost of the haploid deleterious load. Specifically, Kimura (Kimura, 1967) showed that an allele that increases the deleterious mutation rate by Δ*U_d_* bears a fitness cost at equilibrium of *s* = -Δ*U_d_*. Given that deleterious mutations in our simulations are permanently heterozygous and fully recessive we can assume that the deleterious mutation rate in diploid residents is essentially zero. Therefore, Δ*U_d_* is simply the deleterious mutation rate of the haploid and the expected cost of the load in haploids is given by *s* = -*U_d_*. The probability of fixation for a mutation of fitness effect *s* (in a haploid population) is given classically by *P_fix_*(*s, N*) = (1-*e*^-2*s*^)/(1-*e*^-2*Ns*^) (Kimura, 1962). *P_fix_*(*s* = -*U_d_*, *N*) is shown with the red dashed line in Fig. 1A and agrees very well with simulations, confirming that in sufficiently small populations haploid *NP_fix_* is set by the cost of the deleterious load.

Again motivated by our earlier work, we hypothesized that in large populations that favor haploids, haploid *NP_fix_* is elevated above the neutral expectation by selection for beneficial mutations. To confirm the influence of selection for beneficial mutations on haploid *NP_fix_* we assayed the final fitness of populations in which haploids had fixed (Fig. 1B). We found that at smallest *N*, the average fitness of winning haploid populations is, in fact, lower than the ancestral fitness, confirming that at these population sizes, hitchhiking with beneficial mutations (which would necessitate an average fitness of at least 1+*s_b_*) is not a significant influence on *NP_fix_*. Rather, haploids fix in these populations via random genetic drift. However, as Fig. 1B shows, in sufficiently large populations haploid winners’ fitness surpasses 1+*s_b_* for all values of *h_b_* examined. This demonstrates that in large populations haploids fix almost exclusively by hitchhiking. In fact, note that at yet larger *N* haploid fitness may surpass 1+2*s_b_* suggesting that in those populations haploids reach fixation with multiple beneficial mutations. For now, we leave further examination of multiple-mutation (Desai et al., 2007) dynamics in ploidy evolution for future work.

Moreover, we found that at *h_b_* = 0 haploid *NP_fix_* could be well approximated by the expected probability of hitchhiking to fixation with a beneficial mutation, previously derived for a mutator allele by Raynes et al. (2018). In brief, to fix by hitchhiking an invader must first produce a beneficial mutation, which then has to sweep to fixation. The probability that the invader will produce a beneficial mutation is approximately equal to the ratio of *U_b_* to the total mutation rate, *U_b_* + *U_d_*. Since at *h_b_* = 0 the spread of beneficial mutations in the haploid background is unaffected by selection on beneficial mutations in diploids, the probability of the latter for a mutation of fitness effect *s_b_* can be estimated with *P_fix_*(*s_b_, N*) given above. The probability of hitchhiking to fixation is the product of these two probabilities: 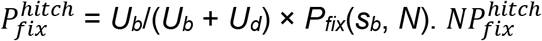 is shown with a teal line in Fig. 1A.

Finally, Fig. 1A also highlights the negative relationship between haploid *NP_fix_* and *h_b_*. As may be expected, *h_b_* has little effect on *NP_fix_* in small populations, in which *NP_fix_* is set by the deleterious load while 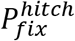 whether for haploids or diploids – is miniscule. However, above *N_crit_* haploids fare worse at higher *h_b_* suggesting that increasing dominance of beneficial mutations lowers 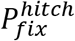 (which dominates *NP_fix_* in at larger *N*). Focusing on the two components of 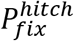 above, we can assume that the probability of a haploid acquiring a beneficial mutation is unaffected by *h_b_*. However, the likelihood of the beneficial mutation escaping drift and reaching fixation may be reduced at higher *h_b_* by stronger competition from beneficial mutations in the diploids. In other words, whereas at *h_b_* = 0, haploids with a beneficial mutation have an advantage *s_b_* over any diploid similarly endowed with a beneficial mutation, that advantage decreases to *s_b_-h_b_s_b_* for any *h_b_* > 0.0. (At *h_b_* = 1.0 haploids enjoy no advantage in the strength of selection on associated beneficial mutations; instead, diploids that benefit from a two-fold higher *U_b_* and a lower deleterious load are favored by selection at all *N*).

To test how interference from beneficial mutations in diploids reduces haploid 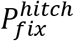 at *h_b_* > 0, we assayed the probability of haploid fixation conditioned on acquiring at least one beneficial mutation, *P*_*fix*|ben_ (Fig. 1C). We found that at *h_b_* = 0, *P*_*fix*|ben_ asymptotes to the expected probability of fixation given by *P_fix_*(*s_b_, N*), which shows that beneficial mutations in haploids are unaffected by competition from fully recessive mutations in diploids. However, at *h_b_* > 0, *P*_*fix*|ben_ declines below the expected *P_fix_*(*s_b_, N*), confirming that competition from beneficial mutations in diploid residents reduces the probability of fixation of new beneficial mutations in haploids. Note also that *P*_*fix*|ben_ declines with increasing *N* and does so faster at higher *h_b_*, consistent with intensifying interference from diploid residents, modulated by both more effective selection and the rising supply of beneficial mutations (i.e., *NU_b_*) in diploids.

Next, we studied the fixation probability of diploid invaders in haploid populations. Once again, we set deleterious mutations to be fully recessive (*h_d_* = 0.0), while beneficial mutations ranged from fully recessive (*h_b_* =0.0) to fully dominant (*h_b_* = 1.0). Fig. 2A confirms that diploids are also susceptible to sign inversion. Unlike that of haploids, though, diploid *NP_fix_* has a negative slope at *N_crit_*. For all studied *h_b_* < 1, diploids are favored in small populations and become disfavored as *N* increases. At *h_b_* = 1.0 diploids are everywhere favored over haploids (have *NP_fix_* > 1 at all *N*).

**Figure 2:**
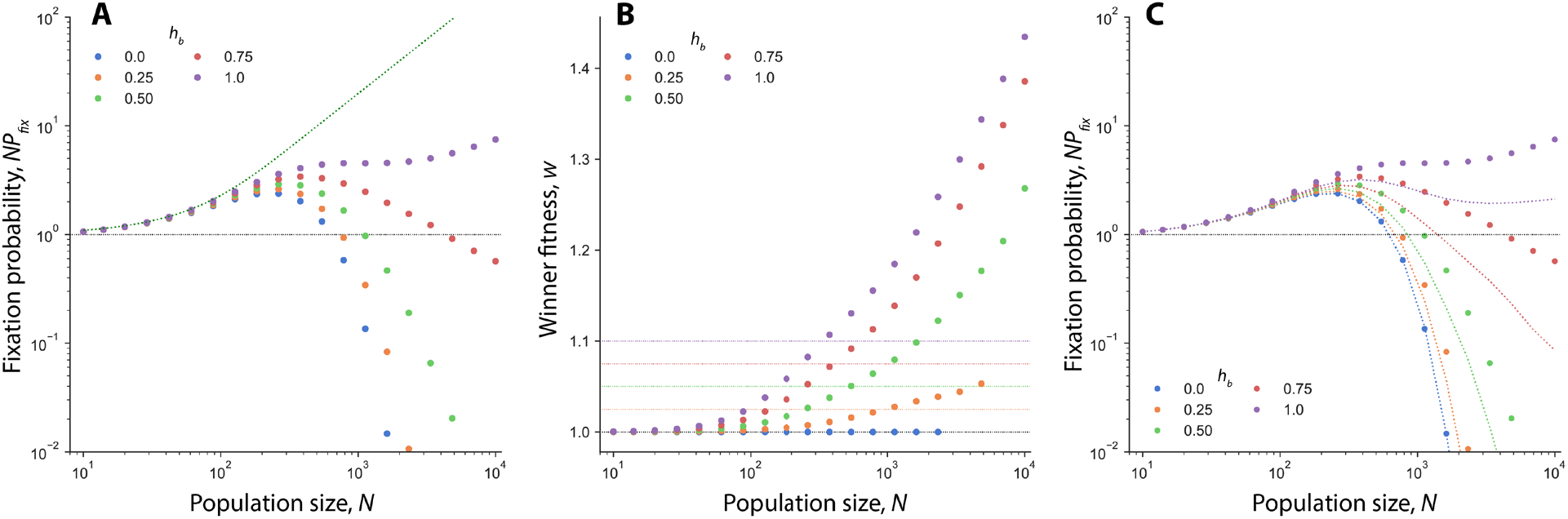
Sign inversion in selection on diploids. **A)** Normalized fixation probability (*NP_fix_*) of diploid invaders (circles) in haploid populations at different *h_b_*. Teal dashed line: *P_fix_*(*s* = *U_d_, N*). Horizontal black line shows the normalized fixation probability of a neutral mutation (*NP_fix_* = 1). **B)** Average fitness of winning diploid populations, *w*. Horizontal lines indicate *w* = 1 in black and *w* = 1 + *h_b_s_b_* in corresponding colors. **C)** Comparison between *NP_fix_* in panel A (circles again) and diploid *NP_fix_* in simulations in which diploid *U* was set to equal that of the haploids (shown for convenience as dashed lines, results only available at the same *N* as circles). Horizontal black line: *NP_fix_* = 1. In all panels, parameter values are as follows: *s_b_*=0.1, *s_d_*=-0.1, *U_d_*= 0.01, *U_b_* = 0.001. All simulation results averaged across 10^7^ replicates.

Our analysis of haploid *NP_fix_* above suggests that in sufficiently small populations, diploids would mostly reach fixation not by acquiring and fixing beneficial mutations but by simply outlasting haploid residents that succumb to selection against the load. Fig. 2A confirms that diploid *NP_fix_* in small populations is well predicted simply by the fixation probability for *s* = +Δ*U_d_* – the diploid’s expected fitness advantage due to the deleterious load borne by haploids. Indeed, assaying the average fitness at the end of simulation confirms that in sufficiently small populations diploid winners are indistinguishable from their ancestors (Fig. 2B).

In larger populations though, in which haploid dynamics are dominated by beneficial mutations, we expected diploid fate to be likewise determined by the likelihood of hitchhiking with beneficial mutations of their own. Correspondingly, we found that the fitness of diploid winners increases with *N* and in sufficiently large populations rises above 1+*h_b_s_b_* (Fig. 2B). This suggests that in those populations diploids also fix exclusively by hitchhiking with beneficial mutations. (Except at *h_b_* =0, where diploid winners never exceed ancestral fitness as hitchhiking with completely recessive beneficial mutations is impossible at any *N*.) For a diploid to fix by hitchhiking, though, a beneficial mutation of effect *h_b_s_b_* must sweep to fixation without interference from beneficial mutations of larger effect *s_b_* in haploids. Diploids possess an inherent advantage of a two-fold higher mutation rate so a diploid invader is always more likely to produce a beneficial mutation than any of its haploid competitors. This advantage is expected to be most pronounced in relatively small populations in which the rate of adaptation is limited by the supply rate of beneficial mutations (Orr & Otto, 1994). Our simulations confirm that the higher *U_b_* improves diploid probability of hitchhiking. Specifically, as shown in Fig. 2C diploid *NP_fix_* is reduced when diploid mutation rate is lowered to that of haploids (see Methods). More importantly, Fig. 2C shows that making diploid *U_b_* equal to that of haploids lowers *N_crit_* for sign inversion for all 0 < *h_b_* < 1.0, suggesting that at some intermediate *N*, diploid *NP_fix_* is above 1 not only due to purifying selection against the load but also positive selection mediated by a higher *U_b_* (note that in simulations without deleterious mutations, diploids are only ever favored by selection at *h_b_* > 0.5 (Fig. S2), consistent with (Orr & Otto, 1994)).

However, as long as *h_b_* < 1, haploids enjoy an advantage in the strength of selection on new beneficial mutations (i.e., beneficial mutations in diploids endow carriers with a smaller fitness advantage *h_b_s_b_* relative to *s_b_* in haploids). This advantage is magnified as population size increases and selection for beneficial mutations becomes more efficient (while the supply rate of such mutations to haploids increases as well). In sufficiently large populations selection for beneficial mutations in haploids offsets the cost of the load and the diploids’ advantage in *U_b_*. When it does, diploid *NP_fix_* (set at that point almost exclusively by diploid 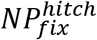, see above) declines below 1 at all *h_b_* < 1.0 (Fig. 2A). Correspondingly, diploids fare better at higher *h_b_*, consistent with a reduced fitness advantage (*s_b_* - *h_b_s_b_*) of beneficial mutations in haploids. And, diploid *NP_fix_* is never below 1 when beneficial mutations are fully dominant: at *h_b_* = 1, haploids have no advantage in the efficacy of selection, whereas diploids still enjoy the benefit of a two-fold higher *U_b_*.

## Discussion

We simulated evolution of ploidy in population with recessive deleterious mutations and beneficial mutations of varying dominance. We showed that in populations with both kinds of mutations, the sign of selection on ploidy can change with population size – a phenomenon we have previously designated sign inversion (Raynes et al., 2018). Consistent with previous expectations (Charlesworth, 1991) for non-equilibrium populations, we found that selection against recessive deleterious mutations lowers haploid fixation probability. Indeed, in sufficiently small populations the evolutionary fate of ploidy variants is entirely dominated by selection against the cost of the deleterious load. Even in the presence of beneficial mutations, haploid invaders that bear the full cost of the deleterious mutations fare worse than the neutral expectation (Fig. 1A). Meanwhile, diploid invaders that mask deleterious mutations and, thus, can avoid the fitness cost of the load fare better than the neutral expectation (Fig. 2A).

On the other hand, in large populations, ploidy evolution is dominated by the indirect effects of selection on beneficial mutations. Previously, it’s been shown that selection for beneficial mutations may favor diploids – that produce twice as many mutations as haploids – when the rate of adaptation is limited by the mutation rate (i.e., the product *NU* is small). We find that the benefit of a 2-fold higher *U_b_* (in tandem with the cost of haploids’ load) may indeed raise diploid *NP_fix_* above the neutral expectation at some *N* that would otherwise favor haploids (Fig. 2C). Conversely, haploids have been predicted to fare better than diploids if new beneficial mutations are sufficiently common that the rate of adaptation is limited not by their supply but by their rate of spread (Orr & Otto, 1994). The reason for this is that mutations in haploids have larger effects on fitness (*s_b_* versus *h_b_s_b_*) and are, thus, more likely to survive drift and reach fixation than mutations in diploids. Our work confirms that in sufficiently large populations the probability of hitchhiking to fixation exceeds the neutral expectation for haploids but is below the neutral expectation for diploids. Correspondingly, in large populations haploids fare better than neutral mutations, while diploid *NP_fix_* drops below 1. The expression for 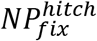 plotted in Fig. 1A for *h_b_*=0 illustrates why haploid *NP_fix_* transitions from below 1 to above 1 with *N*. Recall that 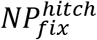 is approximately the product of two probabilities: that of acquiring a beneficial mutation and that of the beneficial mutation reaching fixation. The probability of generating a beneficial mutation (assuming that one is produced) is small and well below one, given that *U_d_* >> *U_b_*. *NP_fix_*(*s_b_, N*) of a beneficial mutation is always above the neutral expectation and is a monotonically increasing function of *N*. And while it may be reduced at *h_b_* > 0 by competition from beneficial mutations in diploids, mutations in haploids should still be favored over those in diploids by a selection coefficient of *s_b_-h_b_s_b_* as long as *h_b_* < 1.0. Thus, when *N* is small, the product of the two, 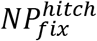, is reduced below the neutral expectation by the low probability of generating a beneficial mutation. However, as population size increases above *N_crit_* and selection on new beneficial mutations becomes more efficient (*NP_fix_*(*s_b_, N*) of new beneficial mutations rises further away from 1), 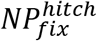 must necessarily increase above the neutral expectation at some sufficiently large *N_crit_*, lifting the total *NP_fix_* above 1 as well.

Taken together, our results demonstrate that sign inversion occurs because ploidy variants can experience both the positive selection mediated by linkage to beneficial mutations and the purifying selection mediated by linkage to deleterious mutations that are overwhelmed by drift at different *N*. That is, haploid *NP_fix_* is dominated by selection on beneficial mutation at large *N* (represented by 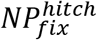) but as population size decreases it is neutralized by drift (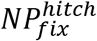 crosses 1) before selection against the cost of the load, which leads to sign inversion.

Previously, we have found evidence of sign inversion in four published studies of the evolution of recombination (Whitlock et al., 2016), cooperation (Nowak et al., 2004), variance in offspring number (Gillespie, 1974), and the mutation rate (Raynes et al., 2018). Having examined all four cases of sign inversion (Raynes et al., 2021) we discovered that all alleles susceptible to sign inversion can impart both fitness costs and fitness benefits on their carriers. That is, similar to ploidy variants, known cases of sign inversion exhibit variability in the sign of selection on individual carriers. We designated such alleles as having sign-variable effects and showed that that all sign-variable effects could be classified into either those that vary between different mutant lineages or among individual members of a single lineage (modulated, for example, by temporal changes in the environment or population composition) (Raynes et al., 2021). Ploidy in our model exhibits between-lineage variability. Haploid lineages that generate beneficial mutations enjoy the indirect fitness benefit and a higher than neutral probability of fixation. Meanwhile, lineages that generate deleterious mutations bear the indirect fitness cost and, correspondingly, have a lower than neutral probability of fixation. More importantly, in the same earlier study we showed that sign variability is necessary for sign inversion, which always occurs because selection on cost-bearing and benefit-endowed carriers exhibit differing sensitivities to drift (Raynes et al., 2021). Our current work showing that sign inversion in selection on ploidy is also driven by between-lineage sign variability corroborates the necessity of sign variability for sign inversion and provides additional support for the common mechanistic basis of sign inversion.

Finally, our findings may be particularly relevant for the growing number of evolution experiments that have documented ploidy transitions in laboratory microbial populations – those expected to be relatively far from equilibrium with access to both beneficial and deleterious mutations. In particular, studies in *Saccharomyces cerevisiae* have repeatedly observed shifts from haploidy to diploidy through spontaneous genome duplication (Aggeli et al., 2021, Venkataram et al., 2016, Fisher et al., 2018, Levy et al., 2015, Voordeckers et al., 2015). Interestingly, most of these studies have been conducted in large populations that, we have found, should indirectly favor haploids rather than diploids. Accordingly, several studies have documented an inherent fitness advantage to diploidy (Aggeli et al., 2021, Venkataram et al., 2016, Fisher et al., 2018) that would allow diploids to fix through the action of direct selection instead. Others, however, did not detect a significant direct advantage to diploid cells over haploids (Dickinson, 2008, Gerstein & Otto, 2011), suggesting that diploids might have arisen via indirect selection. Yet, whereas the appearance of diploids in small populations of the mutation accumulation experiment of Dickinson (Dickinson, 2008) appears consistent with our results, diploids repeatedly overtaking large populations in the study of Gerstein et al. (Gerstein et al., 2006) is difficult to reconcile with our simulations. It is possible that ploidy evolution, at least in yeast, is universally driven by direct selection and an inherent fitness advantage will eventually be identified in all laboratory conditions in which diploids have been shown to overtake haploids (see also (Gerstein & Sharp, 2021) for the discussion of “ploidy drive”). It is also possible, though, that experimental populations differ in some properties that affect indirect selection on ploidy in such a way as to indirectly favor diploids in some environments and disfavor them in others, even at relatively large *N* (see also (Harari et al., 2018)). For example, our work suggests that diploids could fare better in larger populations given a higher *h_b_* or *U_d_* (which would reduce haploid 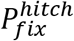), both of which remain somewhat difficult to assay experimentally. All in all, while our work adds to our understanding of the mechanics of indirect selection on ploidy variants, whether indirect selection can be the driving force in ploidy evolution in laboratory or in nature remains to be determined.

