## Supplemental Figures for "Sign inversion in selection on ploidy"

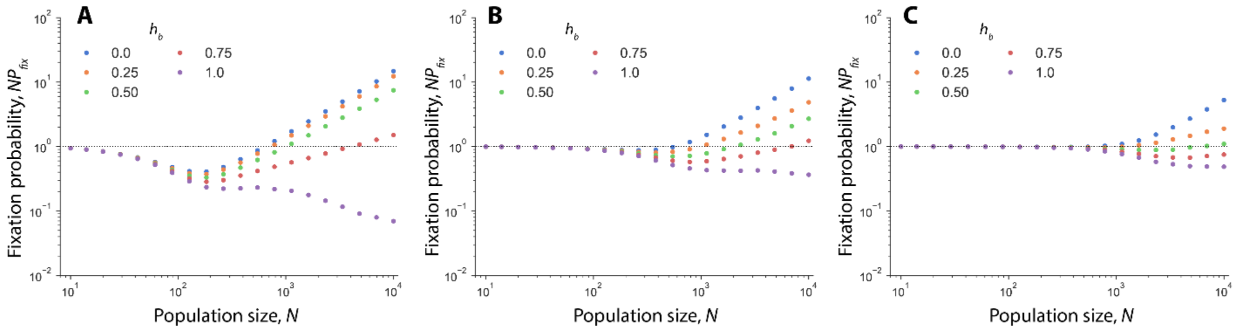


**Figure S1: Haploids are susceptible to sign inversion across a broad range of mutation rates.** Normalized fixation probability ($NP_{fix}$) of haploid invaders in diploid populations at different $h_b$. In all panels $s_b$ = 0.1, $s_d$ = -0.1. Simulation results averaged across $10^7$ replicates. **A)** $U_d$ = 0.01, $U_b$ = 0.001, **B)** $U_d$ = 0.001, $U_b$ = 0.0001, and **C)** $U_d$ = 0.0001, $U_b$ = 0.00001.

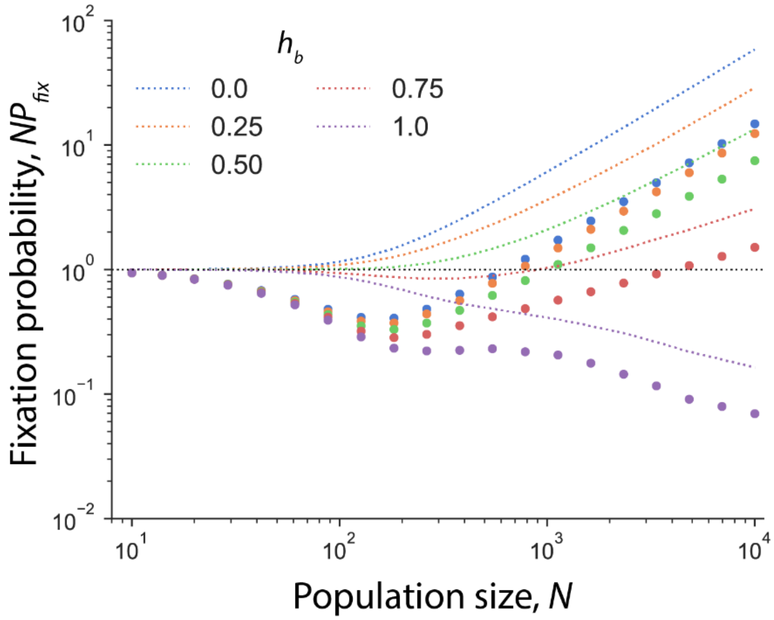


**Figure S2: Haploid $NP_{fix}$ in the absence of deleterious load.** Data from Fig. 1A in the main text are shown again with circles. $NP_{fix}$ at $U_d$ = 0.0 are shown with dashed lines (results only available at the same *N* as circles). $s_b$=0.1, $s_d$=-0.1, $U_d$= 0.01, $U_b$ = 0.001. All simulation results averaged across $10^7$ replicates. Note that at $U_d$=0.0, haploid $NP_{fix}$ is indistinguishable from the neutral threshold in very small populations, consistent with the absence of selection against the load. Furthermore, haploids are disfavored everywhere at $h_b$=1 and at intermediated *N* at $h_b$ > 0.5. This is consistent with the findings of (Orr & Otto, 1994) that diploids may be favored over haploids by virtue of the diploids' advantage in $U_b$, but only in relatively small populations and $h_b$ > 0.5.

Orr, H. A. & Otto, S. P. 1994. Does diploidy increase the rate of adaptation? *Genetics* **136**: 1475-80.
